# The role of gene segment interactions in driving the emergence of dominant gene constellations during influenza virus reassortment

**DOI:** 10.1101/2021.02.10.430697

**Authors:** Sanja Trifkovic, Brad Gilbertson, Emily Fairmaid, Joanna Cobbin, Steven Rockman, Lorena E. Brown

## Abstract

A segmented genome enables influenza virus to undergo reassortment when two viruses infect the same cell. Resulting reassorted progeny have a spectrum of gene constellations and potentially different phenotypes. Although reassortment is involved in the creation of pandemic influenza strains and is routinely used to produce influenza vaccines, our understanding of the factors that drive the emergence of dominant gene constellations during this process is incomplete. Using an influenza vaccine seed production model, reassortant genotypes were tracked through the reassortment process under antibody selective pressure. We discovered that certain gene constellations conferring low replicative fitness were selected at the expense of more fit progeny. Nevertheless, relatively unfit reassortants likely provide high hemagglutinin antigen yields through co-production of non-infectious particles and/or by more hemagglutinin molecules per virion. Our data illustrate the dynamics and complexity of reassortment and highlight how gene segment interactions formed during packaging, in addition to antibody pressure, restrict the final viruses that dominate.

## Introduction

Reassortment, or the swapping of gene segments, is a major mechanism of influenza virus evolution. After co-infection of a single cell by two or more influenza viruses, each viral genome is replicated and various combinations of individual gene segments, in the form of viral ribonucleoprotein complexes (vRNPs), are co-packaged into progeny virions. Since influenza has 8 vRNPs, co-infection with two viruses could theoretically yield up to 256 (2^8^) different gene segment constellations. Although reassortment can occur at high frequency (1-3), studies have shown this to be a non-random process (4-7) with progeny populations restricted due to incompatibilities between gene products (8-10), preferential co-packaging of gene segments (11-13), and certain genes reassorting at higher frequencies (1, 14-16).

Using cross-linking of gene segments *in virio* together with deep sequencing of digested products, we recently uncovered an extensive network of inter-segment interactions believed to be used by the virus to package its genome (17). Contrary to existing dogma, these interactions were not restricted to the previously defined “ packaging sequences” at the ends of the segments but occurred throughout their entire length (11, 17). The pattern of interactions differed markedly between viruses of different subtypes and to a lesser extent between strains of the one subtype (17). Importantly, the observed interactions were numerous, with hundreds being detected in the population of virus particles as a whole. However, some interactions were found at very high frequency, indicating these were likely present in the majority of virus particles. One such high-frequency interaction occurred between the NA and PB1 genes in the early H3N2 virus A/Udorn/307/72 (Udorn) (11, 12) at nucleotides NA 512-550:PB1 2004-2037 (3’ to 5’), equivalent to NA 917-955:PB1 305-338 (5’ to 3’) (17). This NA:PB1 interaction was also maintained in a reverse engineered virus containing the NA and PB1 genes from Udorn and the remaining genes from the H1N1 virus A/Puerto Rico/8/34 (PR8) where the pattern of interactions were found to be essentially inherited from both parent viruses (17). This dominant interaction also likely directed the preferential incorporation of the Udorn PB1 gene during co-infection of Udorn with PR8, even though the resulting reassortant progeny expressing Udorn HA, NA and PB1 were significantly less fit than viruses that contained the Udorn HA and NA genes together with the PB1 gene from the H1N1 strain (11). We proposed that the ability of the influenza virus genome to utilize different sets of interactions to package one of each of the 8 gene segments provides sufficient flexibility to allow reassortment to occur between different influenza viruses, with the stipulation that the stronger interactions are preferentially maintained and shape the gene constellations of resulting reassortant progeny.

Our initial interest in the NA:PB1 gene segment interaction developed from the retrospective analysis of the gene constellations of H3N2 influenza vaccine seed candidate viruses (18). These are produced by reassortment with the highly egg-adapted PR8 virus to enable the creation of viruses displaying greater H3N2 surface antigen yields in eggs through incorporation of non-surface antigen genes from the PR8 parent (19). This classical reassortment process consists of an initial co-infection step in eggs, followed by antibody selection for reassortant viruses with seasonal hemagglutinin (HA) and neuraminidase (NA) surface antigens, and finally cloning by limit dilution to isolate dominant viruses, some of which will display high hemagglutination titres suitable for vaccine seeds. We showed that, unlike other non-surface antigen genes, the PB1 gene of the seasonal H3N2 virus was present in high frequency in H3N2 vaccine seeds and further, that Udorn virus exemplifies this process, with 75% of the progeny virus expressing Udorn PB1 following reassortment with PR8 in eggs (18). Here we address two of the questions raised by these observations. Firstly, we considered what the spectrum of the gene constellations resulting from Udorn: PR8 co-infection was at each stage of the classical reassortment process to better understand the dynamics of co-selection of the seasonal NA and PB1 genes. Secondly, we examined why viruses that display inferior replicative fitness, exemplified by those expressing Udorn HA, NA and PB1 genes, could ever be chosen on the basis of high antigen yield. Our study illustrates the dynamics and complexity of classical reassortment and the impact of gene co-selection in this process. We also present data that highlights the effect of genotype on phenotype in relation to antigen expression.

## Materials and Methods

### Influenza A viruses

The highly egg-adapted A/Puerto Rico/8/34 (PR8; H1N1) virus, currently used as a reassortment partner for the production of H3N2 vaccine seeds, and A/Udorn/307/72 (Udorn; H3N2) as a model seasonal isolate, were used in this study. An eight-plasmid DNA transfection system (20) was used to generate reverse engineered (rg) viruses of specific gene constellations defined using the following standard nomenclature e.g. PR8(Ud-HA,NA,PB1,NP) referring to an isolate containing Udorn HA, NA, PB1 and NP genes with the remaining genes from PR8. All viruses were propagated in 10-day old embryonated hen’s eggs at 35°C for 2 days, then allantoic fluid was harvested and stored at -80°C.

### Classical reassortment

This was performed using a modification of the method devised by Kilbourne (21) (Fig. 1A). Ten-day-old embryonated hen’s eggs were co-infected with PR8 and Udorn viruses (Fig. 1B) and incubated at 35°C in a moist environment for 24 hours. As in seed strain production, different ratios of the two viruses were examined (Fig. 1B). Allantoic fluid was harvested and passaged twice in eggs in the presence of polyclonal anti-PR8 antiserum. For the first antiserum passage, eggs were inoculated with virus-infected allantoic fluid and antiserum added after 1 hour. After 48 hours the allantoic fluid was harvested and genotyped to detect the presence of the Udorn HA and NA genes. Virus-infected allantoic fluid with both Udorn HA and NA genes present was subjected to a second egg passage, this time being pre-incubated with the antiserum for 1 hour prior to inoculation of the mixture into eggs. After 48 hours the allantoic fluid was harvested and genotyped to determine the origin of the HA, NA and M genes. The dominant progeny from classical reassortment were isolated by cloning via limit dilution. Briefly, serial dilutions of the allantoic fluid were inoculated into 5 eggs each and after 48 hours the allantoic fluid was harvested. A hemagglutination assay was performed to detect the highest dilution containing virus for subsequent genotyping. If required, a further round of cloning by limit dilution was performed to obtain a pure population.

**Figure 1.**
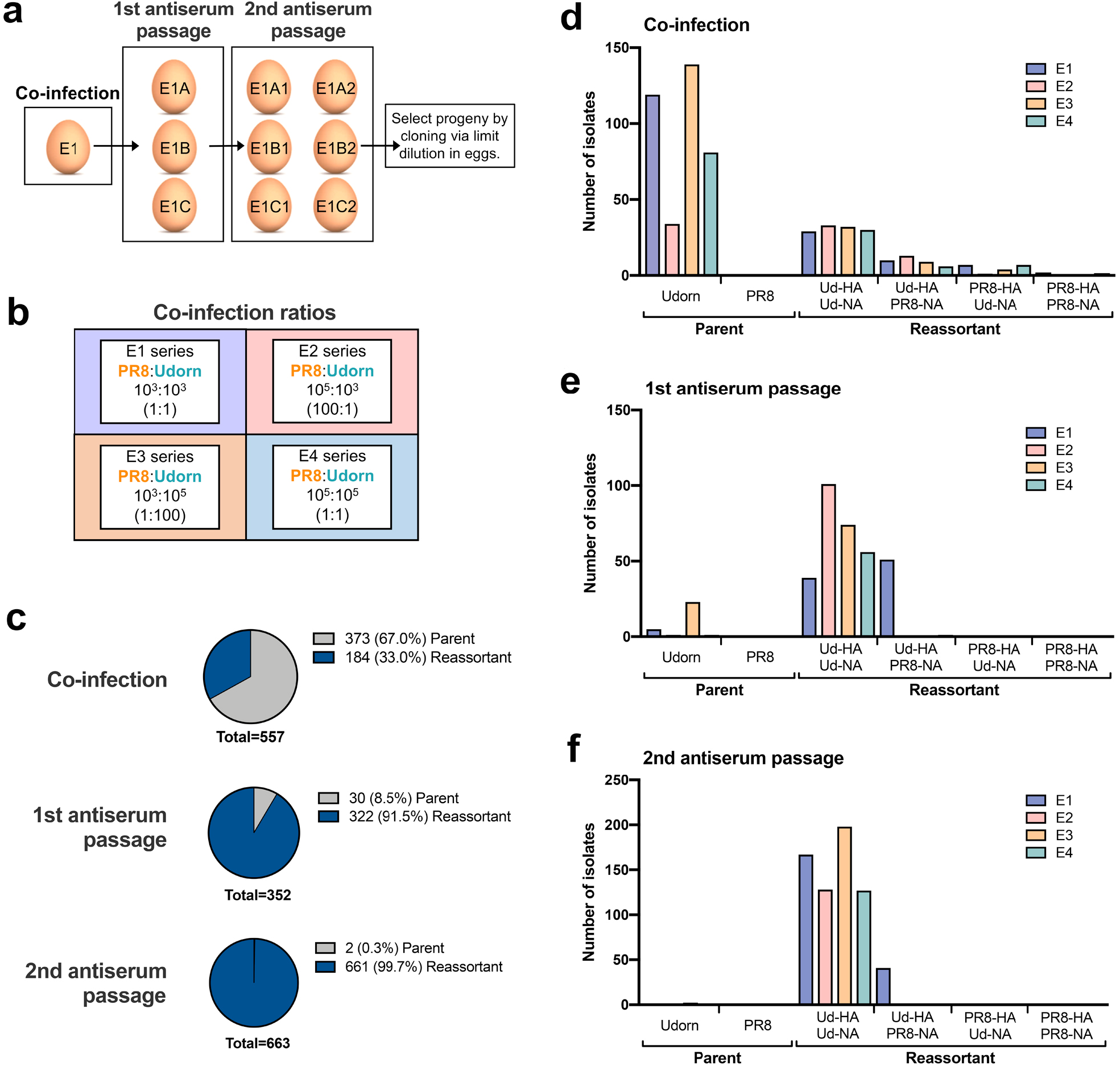
The frequency of parent versus reassortant genotypes isolated throughout the classical reassortment process. **a** Schematic overview of the classical reassortment process used for the generation of seasonal influenza vaccine seeds. Ten-day-old embryonated hen’s eggs were co-infected with PR8 and Udorn at different ratios (series E1-E4). The allantoic fluid from these eggs was then harvested and passaged into 3 eggs (eg. E1A, B and C) in the presence of antiserum against the PR8 surface glycoproteins. The allantoic fluid from each of these eggs was then further passaged into two eggs (eg. E1A1 and E1A2). Viruses from these were cloned by limit dilution. **b** The co-infection ratios and amounts of infectious PR8 and Udorn virus (PFU) used to inoculate the first eggs of each series. **c** The proportion of total parental and reassortant genotypes at each stage of the classical reassortment process. Viruses in the allantoic fluid were isolated by plaque formation. Individual plaques were picked into 0.05% Triton X-100 and gene specific RT-PCR performed to determine the origin of each gene. **d–f** The number of PR8 and Udorn parent genotypes recovered as well as the number of viruses containing different combinations of HA and NA genes present on reassortant viruses in (**d**) the initial coinfection, (**e**) the first antiserum passage and (**f**) the second antiserum passage.

### Virus quantitation

Assays to quantitate hemagglutination (22) (hemagglutinating units, HAU) were performed in microtitre plates with 1% chicken red blood cells and infectious virus yields (plaque-forming units, PFU) were determined by plaque assay on confluent MDCK cells monolayers (23). To establish the relative HA yields of reassortant viruses, a standardised infection was performed by inoculating 10-day embryonated eggs (n=5) with 100 PFU of virus injected into the allantoic cavity. After 2 days at 35°C the eggs were chilled overnight at 4°C before harvesting the allantoic fluid.

### Gene specific RT-PCR for the identification of viral genes

The origin of viral genes present in samples were determined by gene-specific reverse transcription polymerase chain reaction (RT-PCR) using a SensiFast Probe No-ROX one-step reverse transcription kit (Bioline, Meridian Biosciences, Ohio, USA). Each 20 µL reaction contained 5 μl of virus sample in 0.05% Triton-X 100, 10 µL of 2x SensiFast Probe No-ROX one-step master mix, 0.2 µL reverse transcriptase, 0.4 µL RiboSafe RNase inhibitor mix, 0.8 µL of each 10 µM forward and reverse primer and 0.08 µL of each 25 µM gene specific probe. The reverse transcription, amplification and detection were performed in a BioRad CFX96 PCR System. The reaction conditions, primers (Geneworks, Adelaide, South Australia, Australia) and probe (Integrated DNA Technologies [IDT], Coralville, Iowa, USA) sequences are available on request.

### Viral replicative fitness

Viral replication kinetics were determined by infecting MDCK cells using a multiplicity of infection (MOI) of 0.001 PFU/cell. After a 1-hour adsorption period the inoculum was removed, cells were washed and incubated in media supplemented with 1 µg/mL TPCK (designated time = 0 hours). Cell culture supernatants were harvested at various time points and stored at -80°C until required. Infectious viral titres were determined by plaque assay as above.

### Mini genome assay

A β-lactamase (BLA) reporter assay (24) was used to compare the activities of viral polymerase complexes identified in the dominant reassortant viruses as previously described (18). Briefly, 293T cells were transfected with 2 ng each of plasmids expressing the three influenza virus polymerase genes (PB2, PB1 and PA) and the nucleoprotein (NP) gene, together with 2 ng of a plasmid encoding BLA (provided by CSL Ltd.). These pHW2000 plasmids were those used for genetic engineering of influenza viruses in which viral cDNA is inserted into a bicistronic expression system (CMV and RNA polymerase 1 promotors) (20). For the BLA-expressing plasmid the reporter gene was cloned between the 5’ and 3’ non-coding regions of the H1 HA gene to provide specificity for the influenza polymerase complex. Background levels of BLA produced by direct transcription from the CMV promotor was assessed in cells transfected with the reporter gene alone. After incubation at 37°C and 5% CO_2_ for 24 hours, cells were lysed and LyticBLazer™ -FRET B/G substrate (Life Technologies) added. The β-lactamase cleavage of the green substrate to the blue product was measured by optical density every 15 min for a 2-hour period using a CLARIOstar (BMG Labtech, Ortenberg, Germany) fluorescence reader with excitation at 405 nm and blue and green emission detected at 445 nm and 520 nm respectively. Specific β-lactamase activity and thus relative polymerase activity was calculated as follows: (445 nm/520 nm ratio of the sample)/(445 nm/520 nm ratio of BLA plasmid alone).

### Electron microscopy

Transmission electron microscopy was performed at CSL Ltd. by Ross Hamilton using a method modified from that of Hayat and Miller (25). Allantoic fluid was transferred onto Formar-coated copper TEM grids (Athene), which were inverted onto 2% agar plates to remove excess liquid. Grids were negatively stained using 2% sodium phosphotungstate (PTA) pH 7.0, excess stain was blotted away using filter paper (Whatman) and allowed to air-dry. Negatively-stained samples were examined using a Philips CM10 transmission electron microscope running iTEM software (EMSIS GmbH).

### Quantification of viral RNA and viral mRNA/cRNA

To quantitate viral RNA production in MDCK cells, total RNA from infected cells was extracted using an RNeasy Mini Kit (Qiagen). Viral vRNA and mRNA were detected by polarity-specific quantitative RT-PCR using the SensiFast Probe No-ROX one-step kit (Bioline). Each 20 µL reaction contained 5 μl of RNA, 10 µL of 2x SensiFast Probe No-ROX one-step master mix, 0.2 µL Reverse Transcriptase, 0.4 µL RiboSafe RNase inhibitor mix and 0.08 μl of 25 μM gene specific probe. Detection of vRNA or mRNA/cRNA was facilitated by addition of 0.8 μl of 10 μM gene-specific forward or reverse primer respectively. The RT reaction was incubated at 45°C for 10 mins. Following this step, 0.8 μl of 10 μM of the opposite primer was added. In all qRT-PCR assays, serially diluted plasmids of corresponding influenza genes with known copy number were used as standards for quantification.

### Viral protein quantification by slot blot analysis

The relative protein content of virus in allantoic fluid was determined by staining for viral proteins in native conformation using a slot blot apparatus. Viral samples were adsorbed to nitrocellulose using a vacuum pump. The nitrocellulose strips were blocked in PBS with casein for 30 mins at room temperature and then probed for 1 hour with monoclonal antibodies (mAb) recognizing H3 HA (clone 36/2 (26)) or IAV M1 matrix protein (clone MCA401, BioRad). Strips were washed and incubated for 1 hour with a horse radish peroxidase (HRP)-conjugated secondary immunoglobulin (DAKO). The strips were washed and immersed in citrate-EDTA before addition of the TMB substrate for 1 hour. Strips were then washed, dried overnight and digitised. The signal intensities for the viral HA and M1 protein bands were determined using ImageJ analysis software (NIH, Bethesda, Maryland, USA).

### Viral protein quantification by flow cytometry

The relative viral protein content within infected MDCK cells was determined by flow cytometry. Infected MDCK monolayers were harvested 6 hours after a 1-hour absorption period. Cells were fixed, permeabilised and incubated with either clone 36/2 recognising the HA or clone MCA401 recognising M1 for 30 min. Cells were washed and incubated with an Alexa Fluor 488-conjugated rabbit anti-mouse immunoglobulin (Life Technologies) for 30 min. Data were acquired using a FACSCanto II and analyzed with FlowJo X version 10 (Tree Star, Ashland, OR).

### Statistical Analysis

All of the data within this thesis was analysed using GraphPad Prism version 8.4.2 (GraphPad Software Inc., La Jolla, CA, USA). Unless otherwise stated, the data was analysed for statistical significance by either a one-way analysis of variance (ANOVA) with a Tukey post-test or a two-way ANOVA with a Sidak post-test. A probability (p) value of <0.05 was considered to be statistically significant.

## Results

### Classical reassortment generates a diverse range of gene constellations

To examine the spectrum of reassortants generated through the initial stages of the classical reassortment process (Fig. 1a), the model seasonal strain Udorn (H3N2) was reassorted with the highly egg-adapted PR8 strain (H1N1) in four different co-infection conditions representing independent experiments (Fig. 1b). Throughout the process, allantoic fluid from individual eggs was screened to detect hemagglutination and the presence of the Udorn HA and NA genes. Viruses were then isolated by plaque assay and the origin of each gene determined by gene-specific RT-PCR.

After the initial co-infection step, the progeny viruses were isolated by plaque formation in the presence of anti-PR8 antisera, which effectively prevented residual PR8 parent virus or viruses with PR8 surface antigens from dominating the plaque assay (Fig. 1c, d). Of the remaining viruses present after co-infection, cumulative data across the four co-infection conditions revealed that 33% of the viruses were reassortants and 67% were the Udorn parent (Fig. 1c). Consistent with what is observed for a seasonal influenza virus, the Udorn parent was represented least in the E2 series where the co-infection ratio favoured PR8 over Udorn by 100-fold (Fig. 1d). The percentage of reassortants continued to increase throughout the process, with 91.5% of the viruses being reassortants after the first antibody passage and 99.7% after the second antibody passage (Fig. 1c, e, f), indicating strong selective pressure for reassortment.

Genotyping of the progeny viruses showed 63 discrete gene constellations in the initial co-infection and after two additional rounds of replication and antibody selection, this reduced to 39 distinct genotypes (Fig. 2a). Depending on the co-infection ratio and input dose, differences were observed in the gene constellations detected or their frequencies in each reassortment series. In most cases, the dominant reassortant detected in the first and second antiserum passages was not identical. In the E4 series, the dominant reassortant of the second antiserum passage represented between 40 – 53% of the total reassortants isolated and yet only represented approximately 10% of the reassortants in the first antiserum passage. When considering reassortants incorporating the PB1 gene from the Udorn parent (Fig. 2b), it was clear that these dominated after the initial co-infection and after the second round of antiserum selection for the Udorn HA and NA. In the E3 series, where the coinfection ratio favoured Udorn virus, all reassortants isolated after one or two rounds of antiserum selection contained the Udorn PB1 gene. Three reassortant genotypes, PR8(Ud-HA,NA,PB2,PB1,NP,NS), PR8(Ud-HA,NA,PB2,PB1,NP) and PR8(Ud-HA,NA,PB1,NP), all containing Udorn PB1 and NP in addition to the HA and NA genes, were detected in each reassortment series at relatively high frequencies, the proportion of which differed according to the initial co-infection ratio.

**Figure 2.**
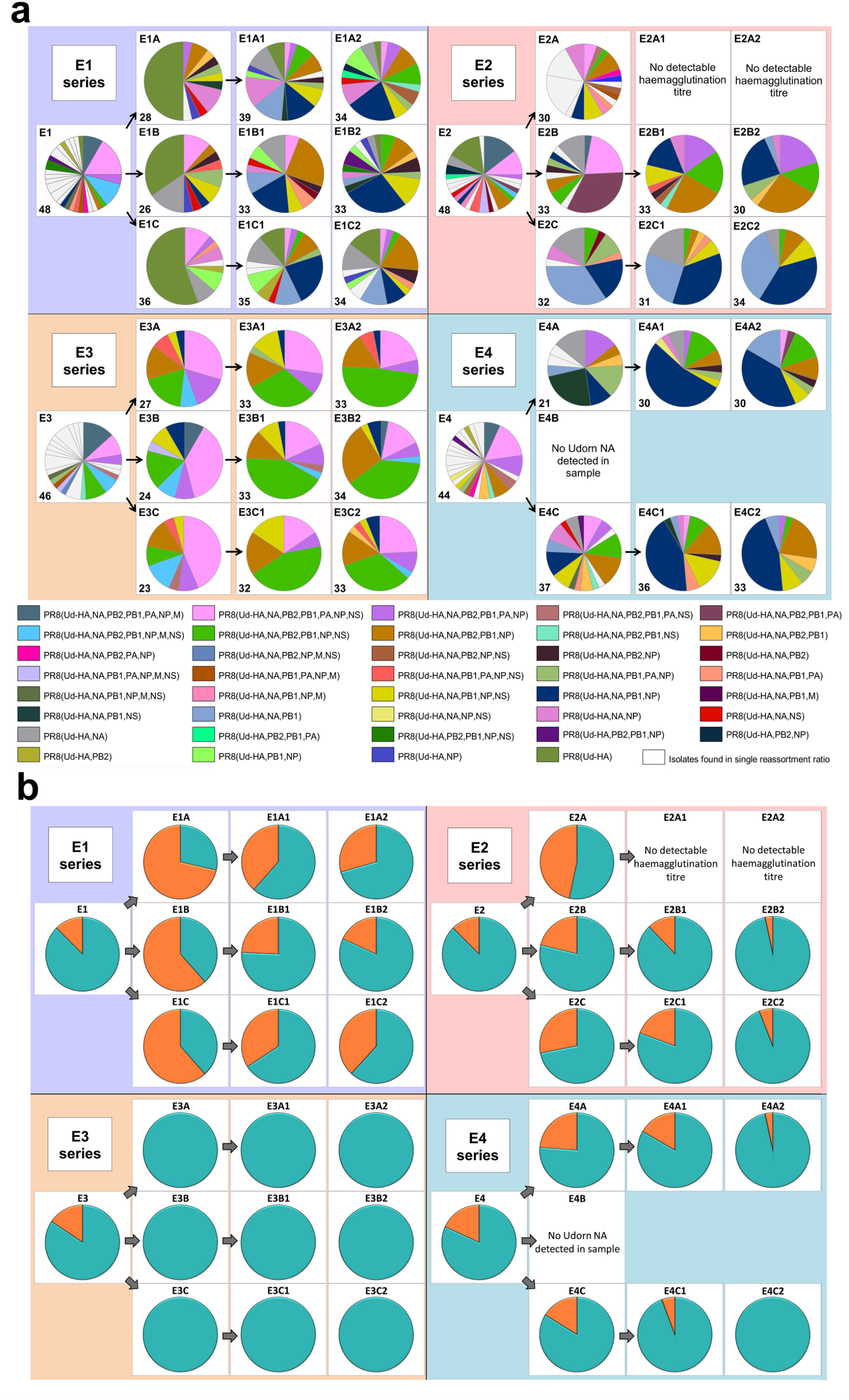
The diversity of reassortant viruses isolated during the reassortment process. Viruses from the allantoic fluid of each egg were isolated by plaque assay. For the co-infected eggs, plaquing was performed in the presence of PR8 antiserum. **a** The genotype of virus in individual plaques was determined using gene specific RT-PCR. Each of the four larger panels represents eggs that originated from the different co-infection ratios (Fig. 1b). The pie charts represent the viruses isolated in the individual eggs corresponding to the co-infection and antiserum passages (Fig. 1a). The dominant gene constellations present in the individual eggs are represented according to the colours in the key. The nomenclature used indicates the Udorn genes on the PR8 background, e.g. PR8(Ud-HA) is a virus with the HA gene from Udorn and the remaining seven genes from PR8. White segments represent gene constellations that were only present in that particular egg. No data is available for eggs E2A1 and E2A2 as the allantoic fluids did not have detectable hemagglutination titres and were not examined further. Initial screening of egg E4B showed that Udorn NA was not present so complete genotypes were not analysed. The number of individual plaques analysed for each egg is indicated in the bottom left of each pie chart panel. **b** The frequency of the Udorn PB1 gene (teal) and PR8 PB1 gene (orange) in the genotypes of the individual virus plaques shown in (a).

### Reassortants that produce high hemagglutination titres in eggs do not always have a high yield of infectious virus

Use of limit dilution to obtain the most abundant reassortants from the second antiserum passage yielded 11 different viruses, 9 of which incorporated Udorn NA and PB1 genes. Notably, PR8(Ud-HA,NA), which is expected to be the high HA yielding vaccine seed virus in our system, was not amongst the dominant genotypes and was made by reverse genetics as a comparator. Eight of the 11 reassortants had significantly higher hemagglutination titres than the parent Udorn virus (Fig. 3a). Four, namely PR8(Ud-HA,NA,PB1), PR8(Ud-HA,NA,PB2,PB1), PR8(Ud-HA,NA,PB1,NP) and PR8(Ud-HA,NA,PB2,PB1,NP) were not significantly different to PR8 virus, illustrating the power of reassortment to enhance the yield of HA from seasonal isolates. To establish whether a greater replicative capacity was responsible for selection of these reassortants, infectious viral titres were determined by plaque assay in MDCK cells (Fig. 3b). Compared to Udorn virus, all but two reassortants had higher infectious virus titres (*p*<0.0001; one-way ANOVA), with six reassortants having equivalent titres to PR8 virus (*p*>0.05; one-way ANOVA). The rgPR8(Ud-HA,NA) virus also had an infectious virus titre that was comparable to PR8 virus (*p*>0.05; one-way ANOVA) and significantly greater than that of Udorn virus and five of the reassortants (*p*<0.0001; one-way ANOVA). These data indicate that the absence of PR8(Ud-HA,NA) virus from the final dominant reassortant progeny was not due to a reduced replicative capacity. Of the four reassortants with hemagglutination titres similar to that of PR8 virus, PR8(Ud-HA,NA,PB1,NP) and PR8(Ud-HA,NA,PB2,PB1,NP) viruses also replicated to the same extent as the PR8 virus (*p*>0.05; one-way ANOVA) yet the two other reassortants, PR8(Ud-HA,NA,PB1) and PR8(Ud-HA,NA,PB2,PB1), did not (P<0.0001; one-way ANOVA). These data indicate that the enhanced hemagglutination phenotype for some reassortants must be due to mechanisms other than the acquisition of greater replication capacity.

**Figure 3.**
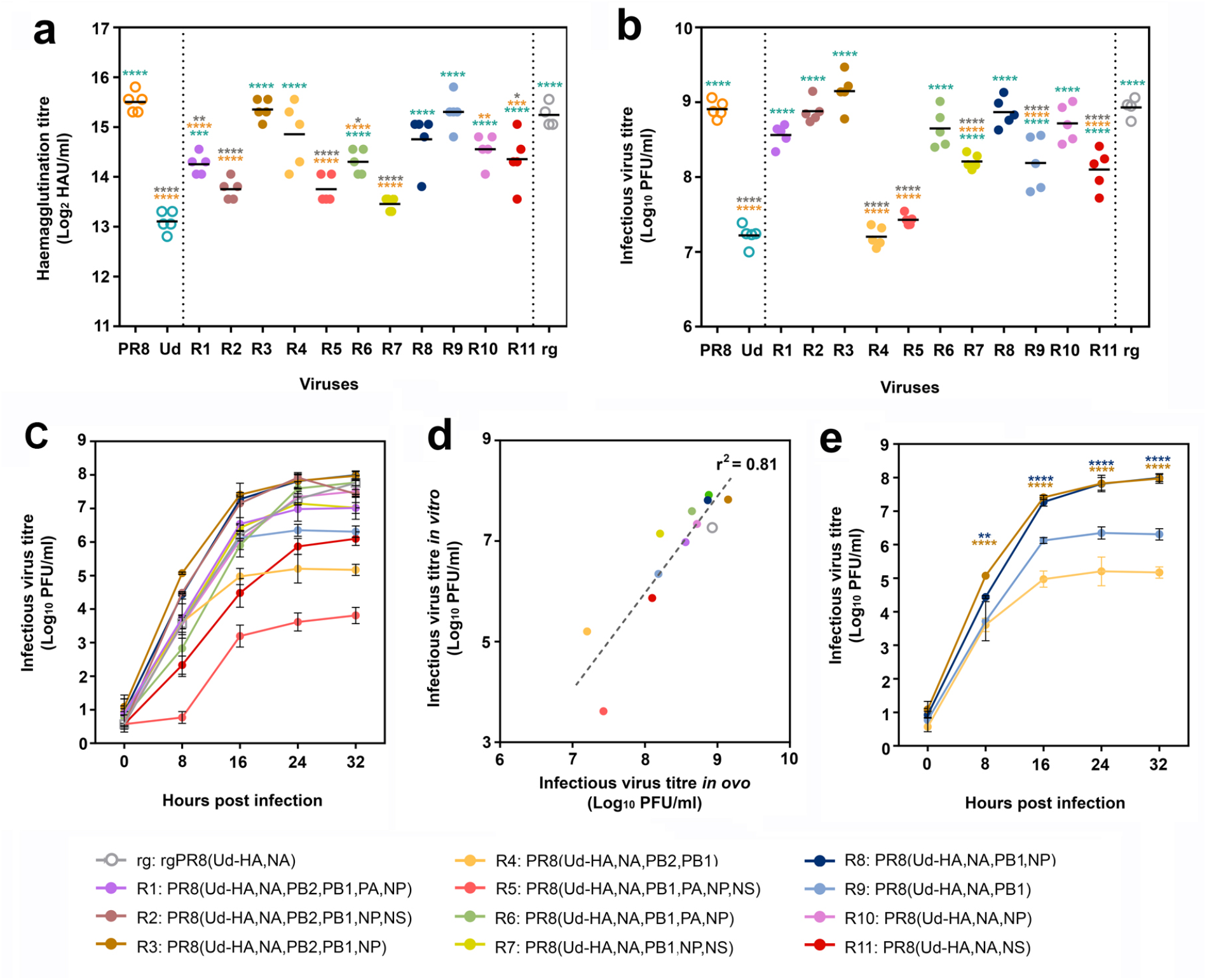
Hemagglutination titres and replicative fitness of dominant reassortant progeny isolated through classical reassortment. **a, b** Eggs were infected with a standard dose (100 PFU) of one of the 11 reassortant viruses (R1-R11) isolated after limit dilution (n = 5), the parental PR8 and Udorn (Ud) viruses (n = 5) or the reverse genetics-derived virus rgPR8(Ud-HA,NA) (n = 4). Virus genotypes are shown in the key by number and colour. Three days post infection, allantoic fluid was harvested and (**a**) hemagglutination titres and (**b**) infectious viral titres determined. Each symbol represents the titre of virus from an individual egg and the line represents the geometric mean. Statistical significance was determined by one-way ANOVA with Tukey’s multiple comparisons test (\**p*<0.05, \*\**p*<0.01, \*\*\**p*<0.001, \*\*\*\**p*<0.0001) for **a** (*F*_13,55_ = 23.39) and **b** (*F*_13,55_ = 61.78). Orange, teal and grey asterisk indicate comparisons of viruses with PR8, Udorn and rgPR8(Ud-HA,NA), respectively. **c** MDCK cells were infected with the panel of reassortant viruses and rgPR8(Ud-HA,NA) at an MOI of 0.001. At the specified time points post infection, virus titres in the supernatants were determined by plaque assay on confluent MDCK cell monolayers. The data represents the mean and standard deviation of three individual experiments. **d** Infectious yields *in vitro* (**c**) at 24 hr as a function of the infectious viral titres *in ovo* (**b**). The dotted line represents a linear regression analysis (*p*<0.0001, r^2^ = 0.81). **e** The data from (**c**) pertaining only to the reassortants with hemagglutination titres equivalent to PR8 *in ovo* (**a**) are shown for clarity. The data represents the mean and standard deviation of three individual experiments. Statistical significance was determined by two-way ANOVA with Sidak’s multiple comparisons test (*F*_3,8_ = 97.28, \*\**p*<0.01, \*\*\*\**p*<0.0001). Brown asterisk indicate comparisons between PR8(Ud-HA,NA,PB2,PB1,NP) with PR8(Ud-HA,NA,PB2,PB1) and blue asterisk indicate comparisons between PR8(Ud-HA,NA,PB1,NP) with PR8(Ud-HA,NA,PB1).

To confirm the phenotypes of the viruses observed in eggs, MDCK monolayers were infected with a standardised dose of the different viruses and viral loads within cultures determined over time (Fig. 3c). Infectious virus yields at the plateau of the replication curve in MDCK cells mirrored the hierarchy found in embryonated eggs as shown by regression analysis of the titres in MDCK cells at 24 hours post infection versus those in eggs (*p*<0.0001, r^2^=0.81; Fig. 3d). Of the four reassortants with hemagglutination titres not significantly different to PR8 virus, PR8(Ud-HA,NA,PB1) had an approximately 1.5 log lower plateau titre than the corresponding virus with the Udorn NP, PR8(Ud-HA,NA,PB1,NP) (*p*>0.0001; two-way ANOVA; Fig. 3e). Likewise, PR8(Ud-HA,NA,PB2,PB1) had a 2.5 log lower titre compared to PR8(Ud-HA,NA,PB2,PB1,NP) (*p*>0.0001; two-way ANOVA; Fig. 3e). Therefore, the addition of the Udorn NP dramatically improved the infectious virus yield obtained when the Udorn HA, NA and PB1 were present in a virion, despite no difference in hemagglutination capacity.

### Reduced replicative capacities of PR8(Ud-HA,NA,PB2,PB1) and PR8(Ud-HA,NA,PB1) viruses are not due to low polymerase activity

It has been proposed that differences in polymerase activity can account for differences in viral replication (9, 27-29). To investigate this, a β-lactamase reporter assay was performed in cells co-transfected with the different combinations of viral polymerase and NP genes represented in the viruses under study (Fig. 4a). Although infection with PR8(Ud-HA,NA,PB1) and PR8(Ud-HA,NA,PB2,PB1) viruses had resulted in lower infectious virus yields compared to the corresponding viruses with Udorn NP (Fig. 3e), their respective RNP complexes (P_PB2_U_PB1_P_PA_P_NP_ and U_PB2_U_PB1_P_PA_P_NP_), demonstrated no significant difference in activity (*p*>0.05; one-way ANOVA; Fig. 4a) to complexes with the Udorn NP (P_PB2_U_PB1_P_PA_U_NP_ and U_PB2_U_PB1_P_PA_U_NP_). Substitution of PR8 PB2 with Udorn PB2 significantly increased the polymerase activity of P_PB2_U_PB1_P_PA_P_NP_ (*p*<0.01, one-way ANOVA) and P_PB2_U_PB1_P_PA_U_NP_ (*p*<0.001, one-way ANOVA), yet no increase in infectious titres was observed for the corresponding viruses both *in vitro* and *in vivo*. These data demonstrate that the viral polymerase activity, as measured by the β-lactamase assay, does not correlate with the efficiency of replication of viruses with these different gene constellations.

**Figure 4.**
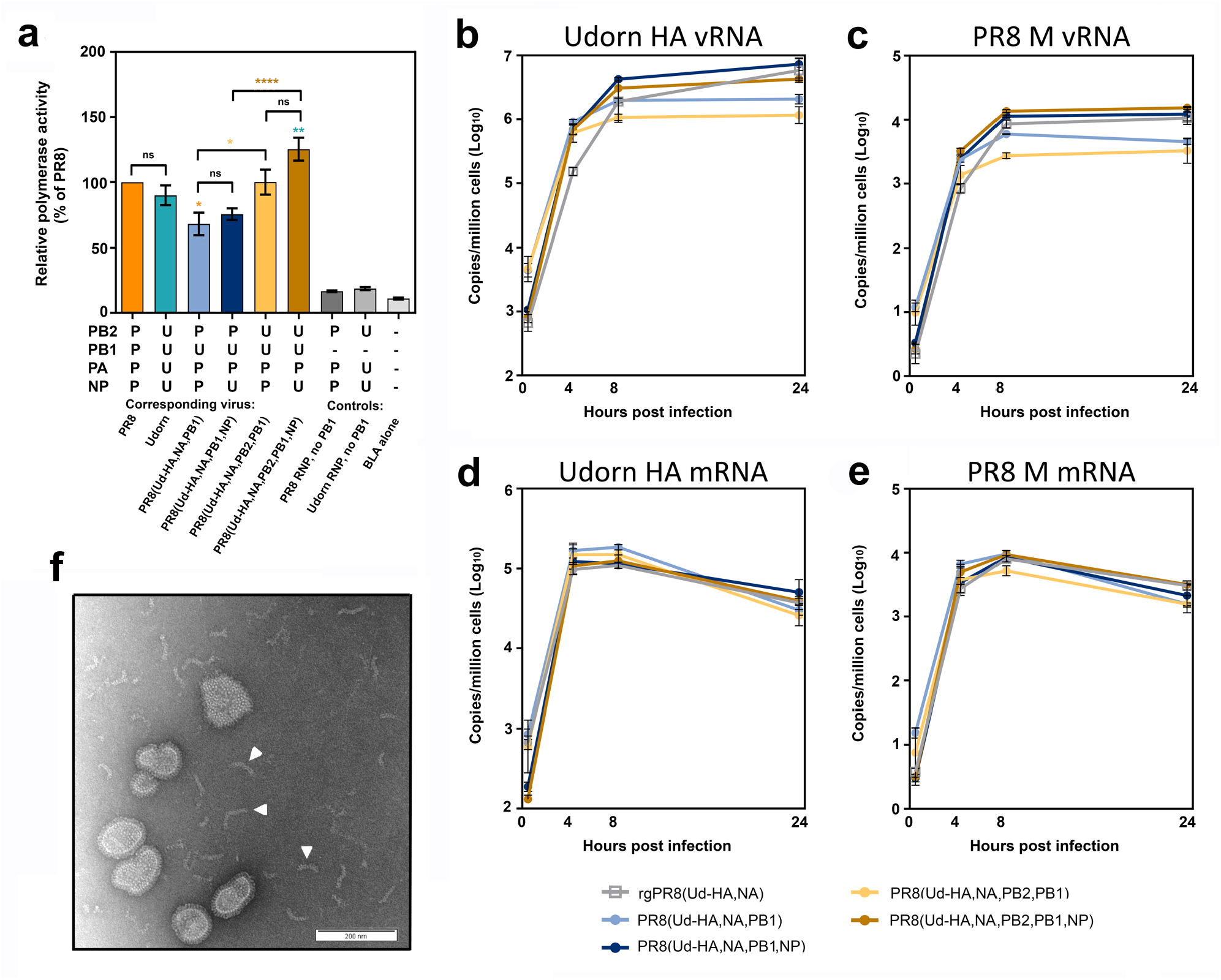
The relative polymerase activity and viral RNA production of the reassortants with high hemagglutination titres. **a** A β-lactamase (BLA) reporter assay was performed in HEK 293T cells that were transfected with the pCAGGS-BLA reporter gene and four pHW2000 plasmids coding for the indicated PB2, PB1, PA and NP genes, which correspond to the RNPs of the indicated viruses or controls without the PB1-encoding plasmid. The relative polymerase activities were normalised to the PR8 RNP activity and each bar represents the mean and standard error of four experiments at the half max of each curve. Statistical significance was determined by one-way ANOVA with Tukey’s multiple comparison test (*F*_8,27_ = 49.28, ns *p*>0.05, \**p*<0.05, \*\**p*<0.01, \*\*\*\**p*<0.0001). Orange, teal, brown and yellow asterisk indicate comparisons between the RNP complexes with P_PB2_P_PB1_P_PA_P_NP_, U_PB2_U_PB1_U_PA_U_NP_, U_PB2_U_PB1_P_PA_U_NP_ and U_PB2_U_PB1_P_PA_P_NP_, respectively. **b-e** MDCK cells were infected with the indicated virus at an MOI of 3 and total RNA extracted from 1×10^6^ cells at 0, 4, 8 and 24 hours after a 1 hour virus absorption period. The copy numbers of (**b**) Udorn HA vRNA, (**c**) PR8 M vRNA, (**d**) Udorn HA mRNA/cRNA and (**e**) PR8 M mRNA/cRNA were assessed by quantitative RT-PCR and the data represents the mean and standard deviation (n = 3) and is representative of four experiments. **f** The presence of free RNP complexes in the allantoic fluid of PR8(Ud-HA,NA,PB1,NP) and PR8(Ud-HA,NA,PB2,PB1,NP) were visualised by negative staining transmission electron microscopy. A representative image is shown with some of the free RNP complexes indicated by a white arrowhead.

### The addition of the Udorn NP to PR8(Ud-HA,NA,PB1) and PR8(Ud-HA,NA,PB2,PB1) results in increased vRNA production

The amount of Udorn HA vRNA (Fig. 4b) and PR8 M vRNA (Fig. 4c), which are common to all the reassortants, were assessed in infected MDCK cells via quantitative RT-PCR. The presence of the Udorn PB1 +/-the Udorn PB2 in a PR8(Ud-HA,NA) background, resulted in significantly reduced levels of HA and M vRNA at 24 hours post-infection compared to the rgPR8(Ud-HA,NA) virus (*p*<0.001 and *p*<0.0001 respectively; two-way ANOVA). Upon the inclusion of the Udorn NP to the PR8(Ud-HA,NA,PB1) and PR8(Ud-HA,NA,PB2,PB1) backgrounds, the HA and M vRNA levels were restored to the levels seen in rgPR8(Ud-HA,NA) at 24 hours post infection. The addition of the Udorn NP to PR8(Ud-HA,NA,PB1) virus increased the amount of HA vRNA at the 8 and 24 hour time points (*p*<0.05, *p*<0.0001 respectively; two-way ANOVA) and also the levels of M vRNA at the 8 and 24 hour time points (*p*<0.01, *p*<0.0001 respectively; two-way ANOVA). Similarly, PR8(Ud-HA,NA,PB2,PB1,NP) displayed significantly higher levels of HA vRNA than PR8(Ud-HA,NA,PB2,PB1) at the 8 and 24 hour time points (*p*<0.001, *p*<0.0001 respectively; two-way ANOVA) and M vRNA at the 4, 8 and 24 hour time points (*p*<0.001, *p*<0.0001, *p*<0.0001 respectively; two-way ANOVA). These data suggest that the higher levels of vRNA produced by viruses with Udorn NP, in addition to Udorn HA, NA, PB1 +/– PB2, may contribute to the greater infectious particle yields of these viruses compared to the corresponding reassortants with PR8 NP. That said, we observed by electron microscopy of the virus-containing allantoic fluid preparations of PR8(Ud-HA,NA,PB1,NP) and PR8(Ud-HA,NA,PB2,PB1,NP) that free RNPs were present in these (Fig. 4d) but not in the corresponding reassortants with PR8 NP, suggesting the possibility that the greater levels of vRNA produced by these Udorn NP-containing viruses may not be all incorporated into virions.

### High hemagglutination titres did not result from increased HA protein in the virion as a result of increased transcription or translation of HA in infected cells

It was possible that the lower levels of HA and M vRNA production observed for PR8(Ud-HA,NA,PB2,PB1) and PR8(Ud-HA,NA,PB1) compared to the corresponding viruses with Udorn NP, might be due to the preferential production of viral mRNA over vRNA. To examine this, the level of Udorn HA and PR8 M viral mRNA transcription was quantified in infected MDCK cells by gene-specific qRT-PCR (Fig. 4d, e). Comparison between rgPR8(Ud-HA,NA) and the four reassortant viruses showed no difference in viral mRNA production for either the Udorn HA or PR8 M genes at any time point (*p*>0.05, two-way ANOVA).

To examine whether selective protein modulation was occurring in our system, the amount of HA present in infected MDCK cells was assessed by flow cytometry 6 hrs post infection (Fig. 5a). Matrix protein was also analysed to determine whether any potential modulation of Udorn HA translation was specific for the HA (Fig. 5b). The data showed a trend towards slightly more HA produced in cells infected with rgPR8(Ud-HA,NA) than with PR8(Ud-HA,NA,PB2,PB1) and PR8(Ud-HA,NA,PB2,PB1,NP) (*p*<0.05; one-way ANOVA). However, the levels of M1 protein expression between infected cells differed considerably with rgPR8(Ud-HA,NA)-infected cells producing more of this protein than the reassortants (*p*<0.0001; one-way ANOVA). When the ratio of HA staining to M1 staining was calculated and expressed relative to rgPR8(Ud-HA,NA) (Fig. 5 key), the fold difference was less than 1.4 suggesting only minor, if any, effects on protein expression specific for HA in infected cells.

**Figure 5.**
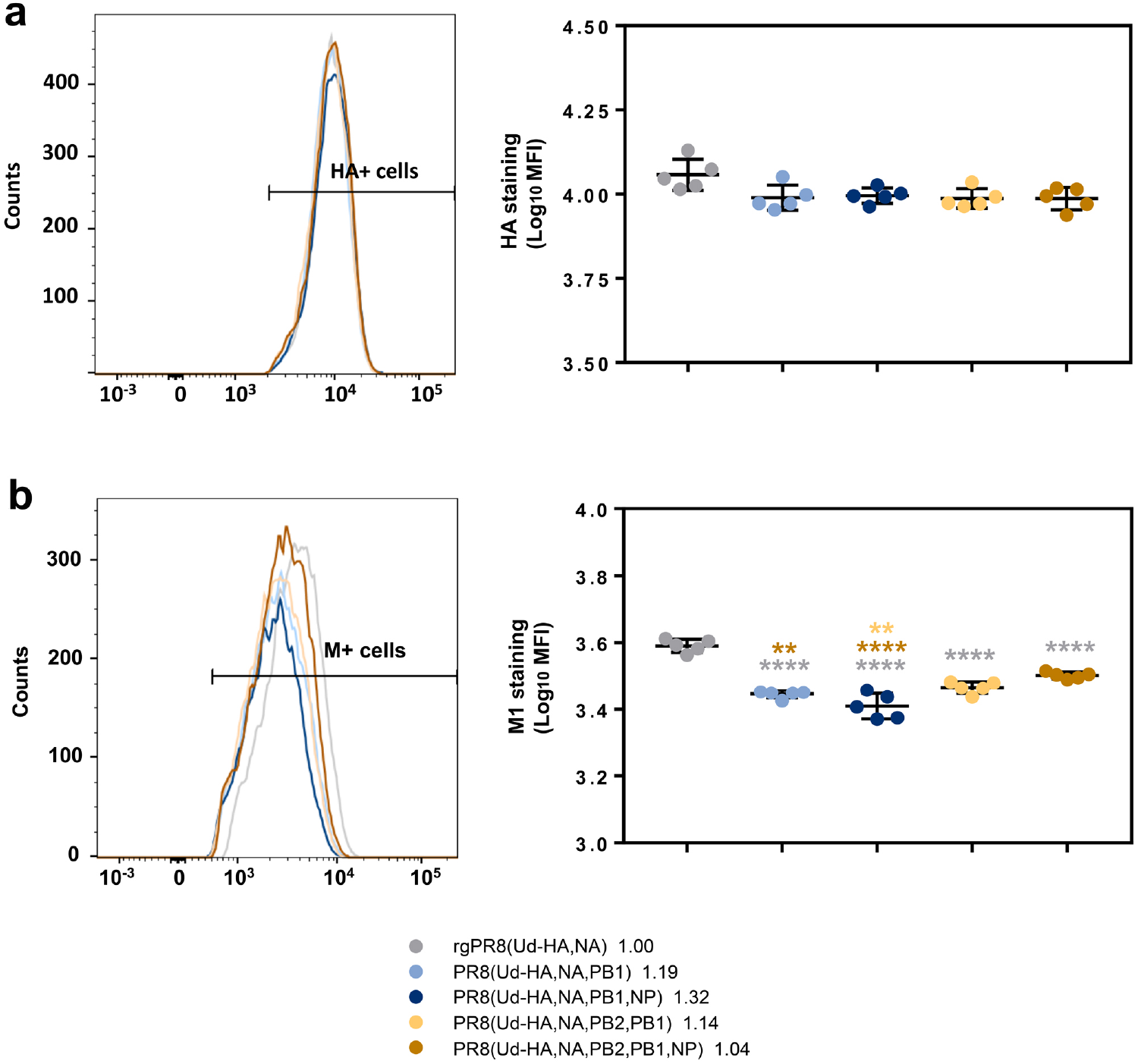
Flow cytometric analysis of Udorn HA and PR8 M1 protein production in infected cells. MDCK cells (1×10^6^) were infected with different viruses at an MOI of 3 and protein production assessed 6 hours after a 1 hour virus absorption period. Cells were stained with either (**a**) anti-Udorn HA (AF647) or (**b**) anti-M1 (MCA401) for the detection of proteins. Left panels show example curves of stained cells; right panels show individual data from 5 replicate cultures, and is representative of 3 experiments. Statistical significance were determined by one-way ANOVA with Sidak’s multiple comparison test (\*\**p*<0.01, \*\*\*\**p*<0.0001) for **a** (*F*_4,20_ = 3.24) and **b** (*F*_4,20_ = 45.59). Grey, brown and yellow asterisk indicate comparison of viruses with rgPR8(Ud-HA,NA), PR8(Ud-HA,NA,PB2,PB1,NP) and PR8(Ud-HA,NA,PB2,PB1) respectively. Values in key represent the ratio of mean HA staining to mean M1 staining.

### An increased number of non-infectious particles and a higher HA density contributes to the high HAU

To further examine what contributes to the high hemagglutination titres of the reassortants we required a measure of relative particle number. Usually this is done by some measurement of the RNA, however, we had observed that the Udorn NP-containing reassortants had free RNPs (Fig. 4f) that would make such measurement misleading. Instead, an analysis of HA protein relative to M1 protein content of the different viruses was performed. As virion size and morphology were similar between the viruses when observed via electron microscopy (Fig. 6a), we made the assumption that the amount of matrix protein (M1) present in the viral particles in the different preparations would be approximately equivalent on average and correlate directly with particle number in the preparation.

**Figure 6.**
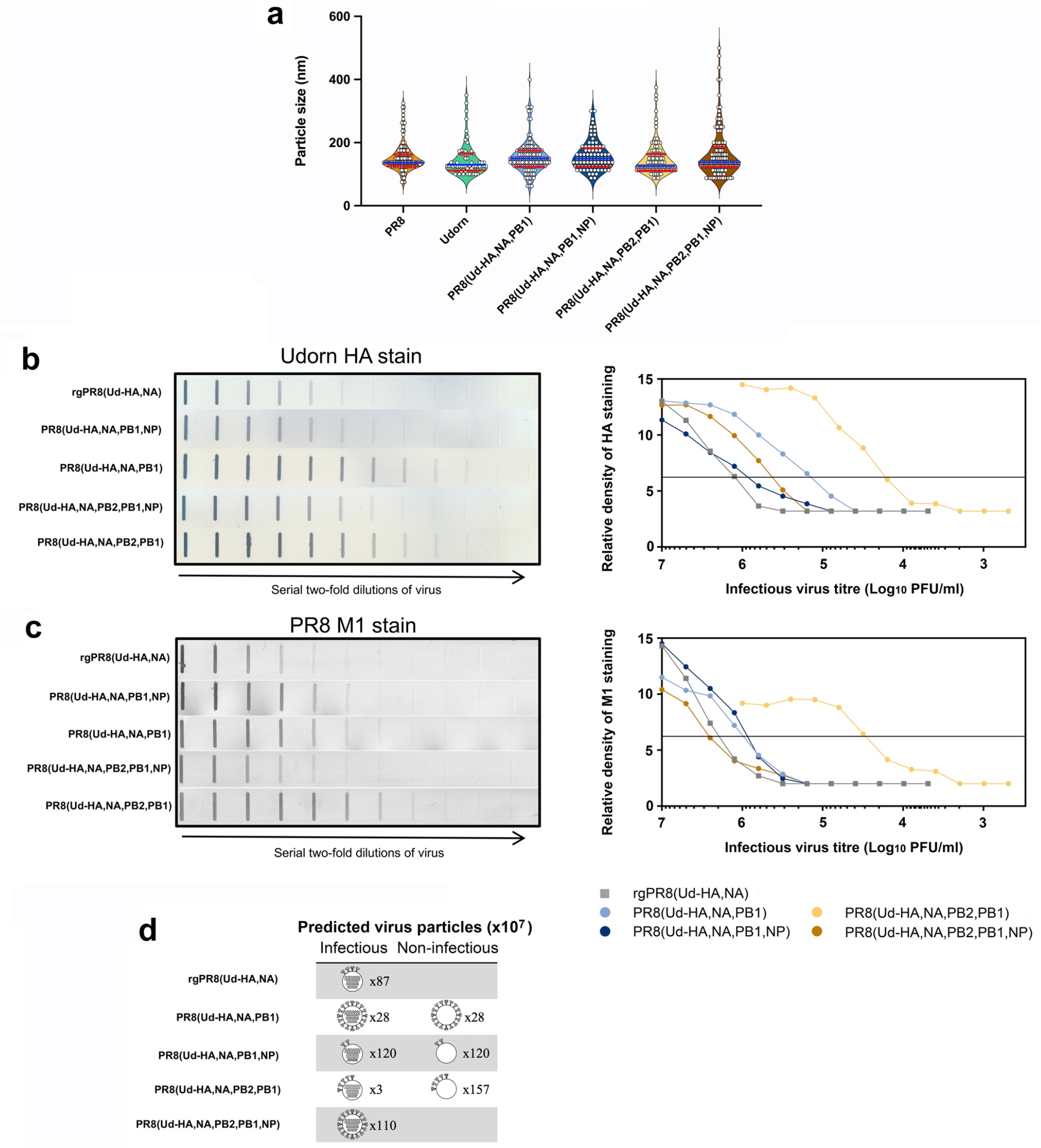
The structure and HA protein content of reassortant viruses. **a** Violin plots of particle sizes visualized by transmission electron microscopy of infected allantoic fluid. White filled circles represent individual measurements along the longest axis of 60-130 distinctive viral particles from one to three printed images per virus preparation converted to nm. The median size is shown as a blue line with quartiles shown as red lines. Analysis by 1-way ANOVA shows no difference in the mean particle size (*p*=0.27) **b, c** Allantoic fluid containing infectious virus was diluted to 10^7^ PFU/mL, or 10^6^ PFU/mL for PR8(Ud-HA,NA,PB2,PB1), and two-fold serial dilutions were performed. Each dilution (100 µl) was transferred to a nitrocellulose membrane and the (**b**) Udorn HA protein or (**c**) PR8 M1 protein was detected with a mouse monoclonal antibody and visualised by secondary staining with rabbit anti-mouse HRP and the addition of substrate. Densitometry analysis was performed on the bands obtained and the peak heights were plotted against the amount of infectious virus in that sample. The horizontal line is used to determine the number of PFU required to provide an arbitrary amount of HA or M1 staining. The data is representative of two experiments. **d** Depiction of the relative amount of HA and its distribution between infectious and non-infectious virions. Infectious particles are depicted with gene segments; non-infectious particles are depicted without gene segments. The predicted number of particles (×10^7^) is indicated beside each virion (Table 2). For ease of representation one arbitrary unit of HA protein (Table 1) is depicted as four HA spikes on the virion. The distribution of the HA protein was assumed to be equal between the infectious and non-infectious particles.

The amounts of HA and M1 in each of the reassortants were determined by the binding of specific monoclonal antibodies using a slot blot assay (Fig. 6b and c). Compared to rgPR8(Ud-HA,NA), less infectious virus was required to provide the same amount of HA staining for each of the reassortants (Fig. 6b), particularly for PR8(Ud-HA,NA,PB2,PB1) where only about one-hundredth of the infectious virus was required (Table 1). With the exception of PR8(Ud-HA,NA,PB2,PB1,NP), less infectious virus was also required to provide the same amount of PR8 M1 staining compared to rgPR8(Ud-HA,NA) (Fig. 6c). By dividing the relative PFU/M1 unit by the relative PFU/HA unit, the resultant HA/M1 ratio indicates the relative amount of HA protein present per arbitrary amount of M1 protein, which, given the assumption of a correlation between M1 and virions, reflects the relative abundance of HA per virion (Table 1). The 3.9-fold greater HA protein density per PR8(Ud-HA,NA,PB1) particle compared to rgPR8(Ud-HA,NA) supports the findings of Cobbin *et al*. (2013) who also identified a 4-fold increase in HA per viral particle for rgPR8(Ud-HA,NA,PB1) by western blotting. The substitution of Udorn NP for PR8 NP in the PR8(Ud-HA,NA,PB1) virus resulted in an 8-fold decrease in HA density from 3.9 to 0.5 HA/M1. The PR8(Ud-HA,NA,PB2,PB1) virus was similar to rgPR8(Ud-HA,NA) with 1.2 HA/M1 and, in this case, the presence of the Udorn NP led to an increase in HA density to 4 HA/M1. Thus, changes in the gene constellation of the virus can dramatically alter the amount of HA on the virion surface.

**Table 1.**
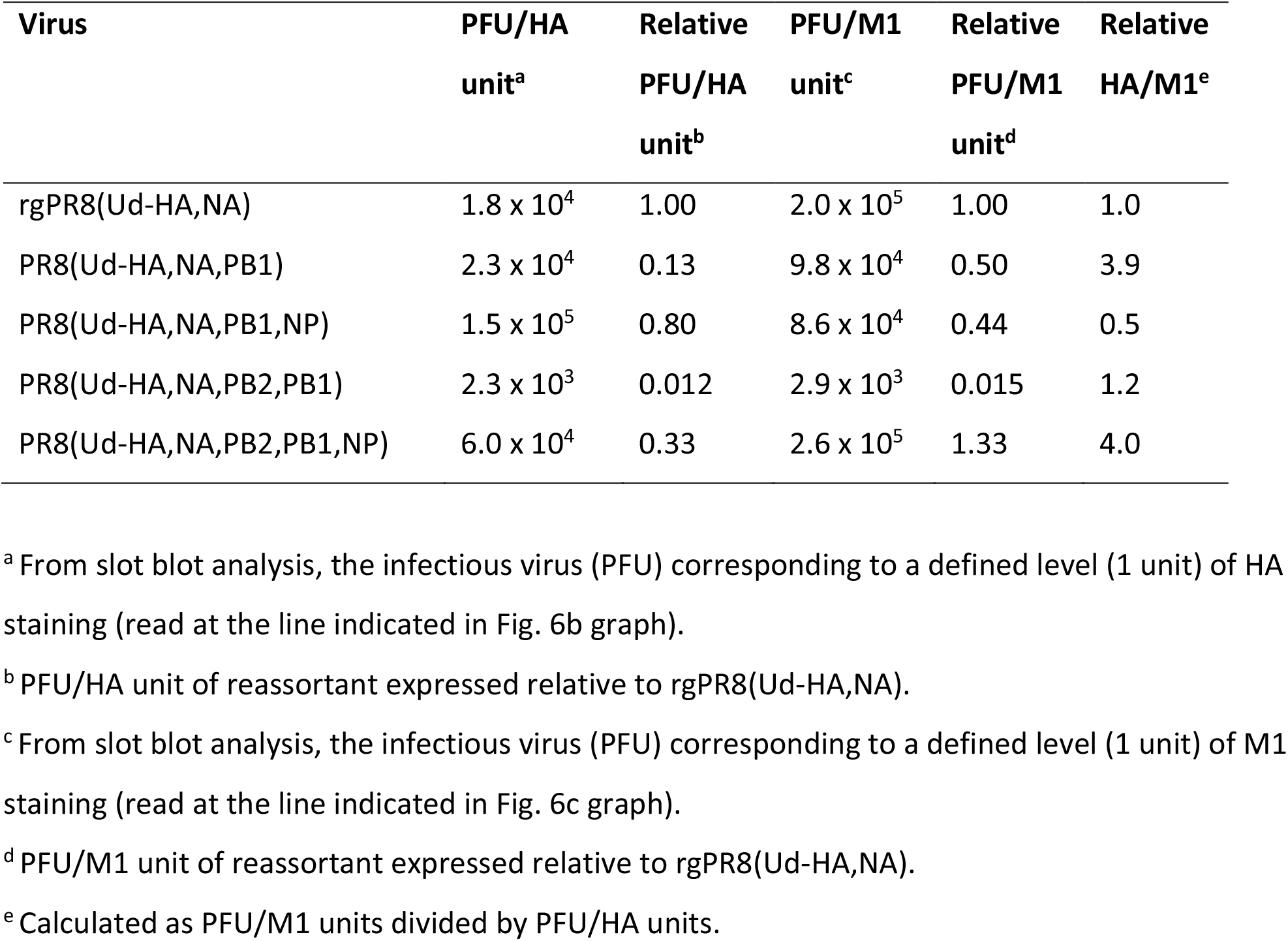
Amount of infectious virus required to achieve an equivalent level of HA or M1 staining and relative abundance of HA per virion.

Data from the slot blot analysis (Fig. 6), taken in conjunction with the infectious virus titres from the allantoic fluid of eggs infected with a standard amount of each of the viruses (Fig. 3b), allows an estimation of how many total virus particles are predicted to be present and the relative total yield of HA from these eggs (Table 2). As an example of these calculations and the underlying assumptions, for a given amount of M1 protein, which we equate to a given amount of virus particles, PR8(Ud-HA,NA,PB1) has only half the number of infectious particles as rgPR8(Ud-HA,NA) (relative PFU/M1 unit Table 1). Assuming for arguments sake that all the virions in the reverse engineered rgPR8(Ud-HA,NA) are infectious, this suggests that only half of the PR8(Ud-HA,NA,PB1) virions are infectious. For a given amount of infectious PR8(Ud-HA,NA,PB1) virions there is an equivalent number of non-infectious virions contributing to the amount of HA in the egg. Dividing the observed titre of infectious virions by the relative PFU/M1 unit ratio for each virus (Table 1) provides the predicted total number of virions in the egg in M1 units/mL. The titre of rgPR8(Ud-HA,NA) from these eggs is on average 8.7×10^8^ PFU/mL (Fig 3b) and assuming these are all infectious, the total particle number is 8.7×10^8^ M1 units/mL. The PR8(Ud-HA,NA,PB1) virus did not replicate as well and showed an infectious virus yield of 2.8×10^8^ PFU/mL, but as this was accompanied by the same number of non-infectious particles (as only 0.5 of the M1 units were infectious), the predicted total number of virions is 5.6×10^8^ virions/mL. As each particle had 3.9 times the amount of HA (Table 1 relative HA/M1), the predicted total yield of HA in the eggs is 2.2×10^9^ HA units/mL or 2.5-fold greater than rgPR8(Ud-HA,NA) (Table 2). Likewise, PR8(Ud-HA,NA,PB1,NP) had a 1.5-fold greater overall yield of HA, PR8(Ud-HA,NA,PB2,PB1) had 2.3-fold and PR8(Ud-HA,NA,PB2,PB1,NP) had 5.3-fold greater HA yield than did rgPR8(Ud-HA,NA) (Table 2).

**Table 2.** Predicted HA distribution on infectious and non-infectious particles

| **Virus** | **Infectious**<br>**virus titre**<br>**(PFU/mL)<sup>a</sup>** | **Predicted total**<br>**virions**<br>**(M1 units/mL)<sup>b</sup>** | **Predicted total**<br>**HA yield**<br>**(HA units/mL)<sup>c</sup>** | **Relative**<br>**HA**<br>**yield<sup>d</sup>** |
| --- | --- | --- | --- | --- |
| rgPR8(Ud-HA,NA) | $8.7 \pm 1.8 \times 10^8$ | $8.7 \pm 1.8 \times 10^8$ | $8.7 \pm 1.8 \times 10^8$ | 1.0 |
| PR8(Ud-HA,NA,PB1) | $2.8 \pm 1.8 \times 10^8$ | $5.6 \pm 3.6 \times 10^8$ | $2.2 \pm 1.4 \times 10^9$ | 2.5 |
| PR8(Ud-HA,NA,PB1,NP) | $1.1 \pm 0.5 \times 10^9$ | $2.4 \pm 0.1 \times 10^9$ | $1.3 \pm 0.6 \times 10^9$ | 1.4 |
| PR8(Ud-HA,NA,PB2,PB1) | $2.4 \pm 0.7 \times 10^7$ | $1.6 \pm 0.5 \times 10^9$ | $2.0 \pm 0.6 \times 10^9$ | 2.2 |
| PR8(Ud-HA,NA,PB2,PB1,NP) | $1.5 \pm 0.2 \times 10^9$ | $1.1 \pm 0.1 \times 10^9$ | $4.6 \pm 0.5 \times 10^9$ | 5.2 |

<sup>a</sup> Infectious titres determined by plaque assay on allantoic fluid from eggs infected with a standard dose of virus (Fig. 3b). Mean ± SD
<sup>b</sup> Observed infectious virus titre (PFU/mL) divided by relative PFU/M1 unit (Table 1). Prediction of total virions assumes all rgPR8(Ud-HA,NA) virions are infectious.
<sup>c</sup> M1 units/mL multiplied by the relative HA/M1 (Table 1). Assumes rgPR8(Ud-HA,NA) has one arbitrary unit of HA per virion.
<sup>d</sup> Predicted yield of HA relative to rgPR8(Ud-HA,NA)

The relationship between HA density per particle, the number of virus particles and the capacity to hemagglutinate chicken erythrocytes is yet to be completely understood. Nevertheless, together these data provide a possible explanation of how high HA yields, predicted from HAU titres, could be achieved by PR8(Ud-HA,NA,PB1) and PR8(Ud-HA,NA,PB2,PB1) despite their poor replicative capacity. This work highlights how different mechanisms operate to achieve high HA yields (Fig 6d) in different reassortants. A higher HA density per virion was the main contributing factor for PR8(Ud-HA,NA,PB1) and PR8(Ud-HA,NA,PB2,PB1,NP) whereas a greater number of non-infectious particles allowed for the high HA yields of PR8(Ud-HA,NA,PB2,PB1) and PR8(Ud-HA,NA,PB1,NP).

## Discussion

In this study we utilised a model of vaccine seed production with the aim of assessing the influence of selective pressures on viral reassortment over multiple passages. Contrary to current thinking, we show that replicative fitness and antibody resistance are not the only determinants dictating which reassortant progeny will come to dominate in this system. We show data compatible with gene co-selection as a powerful force for shaping the viruses that are formed and we document the dynamics of the rise and fall of different constellations across the stages of the process. Though counter intuitive, we show that gene co-selection can lead to less fit viruses being positively selected to the point that they are isolated by limit dilution as dominant progeny. We also asked how these dominant but less fit viruses may display high antigen yields such that they can be chosen as vaccine seed candidates and reveal different phenotypes that are likely to account for this.

Generally, reassortment of H3N2 seasonal viruses with PR8 is thought to increase the yields of HA protein by generating reassortants that replicate to high titres (30) by virtue of their acquisition of a full complement of genes encoding PR8 non-surface antigens. However, from the four reassortants that expressed high hemagglutination titres equivalent to PR8, only PR8(Ud-HA,NA,PB1,NP) and PR8(Ud-HA,NA,PB2,PB1,NP) displayed high infectious virus yields that likely contributed to the high hemagglutination titres achieved. Comparison between the gene constellations of the four final “ vaccine seed candidates” indicated that the presence of the Udorn NP significantly improved infectious virus yields, suggesting some incompatibility of Udorn HA, NA or PB1 with PR8 NP. Viral infectious yields are often associated with the function of the polymerase complex and the relative activity is believed to determine the replicative capacity of a virus (9, 27-29). However, the addition of the Udorn NP had no impact on polymerase activity in the reporter assay, demonstrating that polymerase activity is not necessarily indicative of the replicative ability of a virus. Despite no difference in polymerase activity, in vitro analysis of RNA production demonstrated that viruses with the Udorn NP had significantly increased levels of vRNA, but not viral mRNA, compared to the corresponding viruses with PR8 NP. The NP protein encapsidates vRNA and complementary cRNA but not mRNA and interacts directly with PB1 and PB2 (31). In addition, regions in the NP have been identified as important for vRNA production (32-34), for selective modulation of NA expression (35) and for efficient packaging (35-37). Although virus strain differences in these functions have not yet been investigated, it is possible that a mismatch of NP with PB1 or NA may impair correct packaging of segments resulting in the over-abundance of non-infectious particles in some reassortant genotypes, such as PR8(Ud-HA,NA,PB2,PB1) but not in the corresponding virus with Udorn NP. However it occurs, the restoration of infectious yields by the Udorn NP is possibly contributed to by the increased amounts of vRNA available for packaging into progeny virions to generate a greater number of infectious particles.

In this study, we provide a possible explanation for how dominant gene constellations could provide a high HA yielding phenotype. We show this may be as a result of either a high replicative capacity, the concomitant production of a large number of non-infectious particles, a greater density of HA per virion or a combination of these factors. PR8(Ud-HA,NA,PB1) and PR8(Ud-HA,NA,PB2,PB1,NP) had an overall increase of HA/particle compared to rgPR8(Ud-HA,NA) whereas PR8(Ud-HA,NA,PB1,NP) and PR8(Ud-HA,NA,PB2,PB1) achieved high hemagglutination titres mainly through an increase in the number of total viral particles. Although the calculations used to determine the relative numbers of infectious and non-infectious particles were based on a number of underlying assumptions, the observations on which these assumptions were made, such as infectious yield and the HA and M1 protein content, were all determined experimentally. Therefore, although the actual numbers might be somewhat speculative, the relative differences observed between reassortants are likely to be sound. Previously in our laboratory it was demonstrated that the Udorn PB1 could selectively modulate Udorn HA protein production in cells (18). This as yet undefined mechanism, thought to operate post-transcriptionally (18), may be operating here to explain how PR8(Ud-HA,NA,PB1) and PR8(Ud-HA,NA,PB2,PB1) achieved high hemagglutination titres despite their reduced replication kinetics.

Until recently, viral replicative fitness was considered the main factor to drive the emergence of dominant gene constellations (3, 38). Viruses with lower infectious yields would likely be outcompeted by genotypes with higher replication kinetics, which would then dominate the population. However, in this study, the isolation of six dominant reassortant gene constellations with infectious yields lower than rgPR8(Ud-HA,NA), but not PR8(Ud-HA,NA) itself, demonstrates that viral fitness doesn’t always dictate dominance, at least not in the initial stages of the selection process. Interactions between viral gene segments are known to be important during the assembly and packaging of the eight RNPs into progeny virions (39-41) and the data shown here enforce our previous observation of the preferential co-packaging of the Udorn PB1 and NA gene segments during progeny virion formation (11, 12). We show that the co-selection of the Udorn PB1 gene with the Udorn NA gene is likely responsible for the prevalence of the Udorn PB1 in all stages of the reassortment process and in the final dominant reassortants in our model system. However, we need to entertain the possibility that this co-selection relationship is reinforced by other co-selection relationships between the PR8 genes that might remove them from the available “ packaging pool” . As we are neutralising reassortants expressing PR8 HA and or NA with antisera, genes that co-select strongly with PR8 HA or NA genes will remain unidentified in this study. The fact that the reassortant PR8(Ud-HA,NA) was not isolated as one of the dominant progeny, despite its high replicative capacity and high HA yield, attests to the strength of co-selection in our system and its influence on the availability of genes that would otherwise create highly fit viruses.

The Udorn PB1-NA co-selection relationship is in accord with our retrospective analysis of past H3N2 vaccine seed strains, where the seasonal PB1 was present at a higher frequency compared to the other non-HA and NA genes from the seasonal virus parent (18). The co-selection of internal genes with the HA or NA genes also has direct implications for the generation of pandemic strains. In the event of reassortment between human and avian IAVs, progeny reassortant viruses expressing the surface glycoproteins of the human strain would be inhibited by pre-existing antibodies within the human population, allowing reassortants with the avian HA and potentially also NA, to dominate. As the HA and NA genes may have co-selection relationships with other internal gene segments, this can dictate which avian internal genes are also carried through into the human-infecting strain, shaping the phenotype of the emergent virus and influencing its impact on the human population. For example, co-selection of an avian PB1 would allow expression of a full-length and inflammatory PB1-F2 (42, 43) which has been shown to be a driver of severe secondary bacterial infection (44).

Our study provides new information on the drivers of influenza virus reassortment and the factors that may influence the phenotype of dominant progeny. The eventual move from classical reassortment to reverse engineering for vaccine seed generation will require this understanding so that gene constellations that provide greatest HA and NA protein content can be produced. In addition, a greater understanding of the factors that dictate gene constellations and their corresponding phenotypes likely to arise by reassortment between influenza viruses of human and other reservoir species, will help in prediction of the likelihood and impact of future pandemics.

## Acknowledgements

The work was supported by a National Health and Medical Research Council of Australia Program grant ID1071916 to LEB. ST was supported by an Australian Postgraduate Award. We thank Ross Hamilton from CSL Ltd for providing the electron microscopic images.

## Author contributions

ST performed all experiments with the exception of the hemagglutination assays, which were performed by EM, and genotyping of certain limit dilution viruses by JC. ST analysed the experiments and wrote the draft manuscript. LEB, BG and SR supervised the work, further analysed the data and contributed to the writing of the submitted manuscript.

